# Mitochondrial ATP-Sensitive K^+^ Channels (MitoK_ATP_) Regulate Brown Adipocyte Differentiation and Metabolism

**DOI:** 10.1101/2025.01.21.634060

**Authors:** Osvaldo R. Pereira, Julian D.C. Serna, Camille C. Caldeira da Silva, Henrique Camara, Sean D. Kodani, William T. Festuccia, Yu-Hua Tseng, Alicia J. Kowaltowski

## Abstract

Brown adipose tissue (BAT) plays a central role in mammalian non-shivering thermogenesis, dissipating mitochondrial membrane potentials through the activity of uncoupling protein UCP1 to release heat. Inner membranes of mitochondria are known to be permeable to potassium ions (K^+^), which enter the matrix either through ATP-sensitive channels (MitoKATP) or leakage across the bilayer driven by inner membrane potentials. Mitochondrial K^+^ influx is associated with increased osmotic pressure, promoting water influx and increasing matrix volume. Since BAT mitochondria have lower inner membrane potentials due to uncoupling protein 1 (UCP1) activity, we hypothesized this could involve compensatory changes in MitoKATP activity, and thus tested MitoKATP involvement in brown adipocyte activities under basal and stimulated conditions. We find that cold exposure and adrenergic stimulation in mice modulate BAT MitoK levels, the channel portion of MitoKATP. Genetic ablation of the gene that codes for the pore-forming subunit of MitoKATP in human pre-adipocytes decreased cellular respiration and proliferation, compromising differentiation into mature adipocytes. In mouse cell lines, the absence of the protein limited cellular oxygen consumption in the precursor stage, but not in mature adipocytes. Interestingly, inhibition of MitoKATP in mature adipocytes increased adrenergic-stimulated oxygen consumption, indicating that shutdown of this pathway is important for full BAT thermogenesis. Similarly, MitoKATP inhibition increased oxygen consumption in BAT mitochondria isolated from mice treated with beta 3 adrenergic receptor agonist CL316,243. Overall, our results suggest that the activity of MitoKATP regulates differentiation and metabolism of brown adipocytes, impacting on thermogenesis.

**New and Noteworthy:** Brown fat cells are important to maintain healthy body weight by promoting mitochondrial uncoupling. Here, we demonstrate that mitochondrial ATP-sensitive potassium channels (MitoKATP) have important roles both in the differentiation of brown fat cells and in the activation of energy-dissipating uncoupling in this tissue.

## Introduction

The brown adipose tissue (BAT) is essential for thermoregulation in mammals, with a central role in non-shivering thermogenesis (1–3). Heat production in BAT is accomplished through the activity of uncoupling protein 1 (UCP1), an inner mitochondrial membrane protein that dissipates the proton gradient generated by the electron transport chain, converting energy from substrates into heat, instead of ATP (4, 5). To support high electron transfer rates in mitochondria and UCP1-mediated thermogenesis, BAT oxidizes glucose, free fatty acids, and triglycerides as substrates for thermogenesis (6).

In addition to its importance in thermal homeostasis, BAT-induced heat generation is also highly energy-demanding (7), and therefore can act as an energy sink, counterbalancing nutrient overload. As a result, the tissue has gained a large amount of attention in recent years for its potential therapeutic value under conditions such as obesity (5–10). In this sense, understanding the mechanisms underlying BAT activation could uncover new mechanistic approaches on how to regulate nutritional homeostasis.

Mitochondria in most tissues exhibit high inner membrane potentials (ΔΨ), which leads to marked K^+^ leak into the matrix, given the attraction of cations to the negatively charged matrix and the abundance of K^+^ in the cytosol. Indeed, Peter Mitchell predicted the need for the presence of a K^+^/H^+^ exchanger in the inner mitochondrial membrane when developing the chemiosmotic theory (13), due to the need to continuously counteract the predicted K^+^ leak (14). In excess, K^+^ entry into the mitochondrial matrix promotes water transport, which could lead to matrix swelling and loss of organellar integrity (14–16). Leucine zipper-EF-hand containing transmembrane protein 1 (LETM1) is an inner membrane protein with a crucial role in K^+^/H^+^ exchange activity, but this pathway is not specific, and studies involving LETM1 disruption also show loss of other monovalent cation homeostasis (17).

Interestingly, mitochondria also present a regulated, ATP-inhibited, entry pathway for K^+^ ions into mitochondria, the ATP-sensitive mitochondrial potassium channel (MitoKATP), recently molecularly identified (18, 19). The uniporter is composed of a mitochondrial inner rectifier K^+^ channel (MitoK, Ccdc51) and a regulatory subunit (MitoSUR, Abcb8) (19). MitoSUR is inhibited by ATP, ADP, and selectively pharmacologically activated by diazoxide (16), an effect that can be reversed by and sulphonylureas such as glibenclamide, which, interestingly, do not inhibit the channel in the absence of diazoxide or other agonists (20). Activation increases mitochondrial matrix volumes (15), promotes cristae remodeling (21), regulates mitochondrial redox balance (22–25) and protects tissues against ischemic damage (26, 27).

The decrease in BAT mitochondrial membrane potentials associated with thermogenesis should reduce rates of mitochondrial K^+^ leak, as this process is highly reliant on differences in charge accumulation as its driving force. Because of this, we hypothesized that MitoKATP-mediated K^+^ entry into mitochondria in this tissue may be particularly important to maintain matrix volume homeostasis in the absence of high inner membrane potentials. We designed this study to uncover the role of MitoKATP channels in BAT function, and surprisingly found that it is involved in BAT differentiation. Furthermore, we found that modulation of MitoKATP activity is important for full thermogenic activation of BAT.

## Methods

### Animals

Male C57BL/6J mice were obtained from the *Biotério do Instituto de Ciências Biomédicas, Universidade de São Paulo*. C57BL/6N were obtained from the *Biotério da Faculdade de Medicina, Universidade de São Paulo*. All experiments were conducted in agreement with the NIH Guidelines for the humane treatment of animals and were approved by the local Animal Care and Use Committees (20/214 and 6071181120). For cold versus thermoneutrality exposure, 8-week-old male C57BL/6J mice were kept either at 10 or 30°C (± 1°C) for 14 days in an environmental chamber (NQ1 model, Environmental Growth Chamber, Chagrin Falls, OH) (28). After this period, mice were euthanized by ketamine and xylazine overdose (300 mg/kg and 30 mg/kg, respectively). For cold exposure experiments, animals were kept in individual cages to avoid group heating behavior. For CL316,243 (CL316) treatment, 8-week-old C57BL/6N male mice were kept under standard housing conditions. Animals received daily IP injections of 1 mg/kg CL316,243, or equivalent volumes of saline for 14 days (29–31). Body weight and food intake were measured every 3 days during treatment. Euthanasia was carried out as described above.

### Cell Cultures

9B [immortalized murine brown preadipocytes, (32)], WT-1 [immortalized murine brown preadipocytes (33)], and A41hBAT cells [immortalized human brown preadipocytes (34)] were cultivated in DMEM (25 mM glucose, and 2 mM glutamine, 11965092, Thermo Fisher Scientific, USA) supplemented with 10% fetal bovine serum, 100 U/ml penicillin and 1000 mg/ml streptomycin, and kept in 37°C incubators with 5% CO_2_. Two days after reaching confluence, WT-1 mouse cells were differentiated by supplementing with 500 μM IBMX, 1 nM T3, 125 μM indomethacin, 5 μM dexamethasone, and 20 nM insulin in the culture media for 6 days, changing media every 2 days, as previously described (35, 36). Similarly, human cells were grown until confluence, and, after 2 days, adipogenesis was induced by supplementing 0.1 μM insulin, 2 nM T3, 500 μM IBMX, 30 μM indomethacin, 0.1 μM dexamethasone, 33 μM biotin, and 17 μM pantothenate for 19-22 days, as previously described (37, 38).

### Primary Adipocytes

4-week-old male C57BL/6N mice kept under standard housing conditions were euthanized as described above. Under sterile conditions, BAT was dissected from three to six animals and maintained in culture media (high glucose DMEM supplemented with 10% newborn calf serum, 10 mM HEPES, 4 mM glutamine, 100 IU penicillin/streptomycin and 25 μg/mL ascorbate). After dissection, the tissue was suspended in digestion buffer (123 mM NaCl, 5 mM KCl, 1.3 mM CaCl_2_, 5 mM glucose, 1.5% FFA BSA, and 100 mM HEPES, pH = 7.4 NaOH, filtered sterile), cut with scissors and digested for 35 min at 37°C, vortexing every 5 min. The resulting suspension was homogenized and filtered through a 100 μm cell strainer. The filtered fraction was kept over ice for 30 min to separate the superior fat layer from the cell debris pellet. The infranatant containing preadipocytes was collected and filtered through a 40 μm cell strainer. Cells were pelleted by 700 g centrifugation for 10 min, washed with DMEM and centrifuged once more. The final cell pellet was suspended in culture media and plated for experiments. Adipocyte precursor cells were differentiated in culture media supplemented with IBMX, T3, indomethacin, dexamethasone, and insulin for 7 days. Differentiation was confirmed by observing lipid droplet accumulation using an optical microscope (39).

### Generation of Crispr-Induced MitoK Mutant Cell Lines

Genomic sequences from Homo Sapiens and Mus musculus MitoK (Ccdc51) were obtained from the UCSC Genome Browser (40). Exon 3 was used as input to obtain gRNA sequences, which are listed in Table 1, using the CRISPOR tool (41). LentiCRISPR v2 (Addgene #53961) and lentiCRISPR v2-neo (Addgene #98292) were kind gifts from Feng Zhang and Brett Stringer, respectively. gRNA sequences (gNT TCTGATAGCGTAGGAGTGAT, Hs g1 TGGACTCAACGAGGTTCGAG, Hs g2 AGAGTTTGTTGGACTCAACG, Mm g1 CGTGCTGCATTGCGAACACG, Mm g2 GTTCTCCGTACCAGTATACG) were cloned into plasmid backbones using BsmBI restriction sites (42, 43). Insertion was confirmed by Sanger sequencing. LentiX 293T cells were co-transfected with 5 μg LentiCRISPR as well as 3.75 μg psPAX2 and 1.5 μg pMD2.G, kind gifts from Didier Trono (Addgene # 12260 and # 12259). Media obtained from LentiX 293T cells was centrifuged at 600 x g for 5 min and filtered through a 0.45 μm nylon filter. 1 mL of each lentivirus-containing media supplemented with polybrene (10 μg/mL) was used to infect 40.000 preadipocyte cells. WT-1 cells were selected with 500 μg/mL geneticin (ThermoFisher Scientific, USA), and A41hBAT cells were selected with 10 μg/mL puromycin (InvivoGen, USA) for 1 week. As a control, parental cells were also challenged with antibiotics to ensure that treatment was effective to completely kill non-resistant cells.

### Generation of shRNA MitoK KD Cell Lines

pLKO.1 neo (Addgene #13425) was a kind gift from Sheila Stewart. A shRNA sequence targeting the mouse Ccdc51 was obtained from the RNAi Consortium Library through Sigma’s website. The targeting sequence (GAAGAGAAGAGGCTCCGAATA), as well as a scramble sequence (CCTAAGGTTAAGTCGCCCTCG), were cloned into pLKO.1 neo using AgeI and EcoRI restriction sites (44). Insertions were confirmed by Sanger sequencing. Lentiviruses were produced through the same protocol described for LentiCrispr, and viruses were used to infect preadipocytes, which were then selected as described.

### Proliferation Assay

5.000 A41hBAT cells/well were seeded at day 0 in four 24-well plates. Cells were fixed with 10% buffered formalin for 15 min at room temperature on the same day, as well as in the following ones (days 2, 4, and 6). Cells were washed and kept in PBS at 4°C until the final day of the experiment. PBS was removed and fixed cells were stained with crystal violet solution (2 mg/mL, 20% MetOH) for 15 min at room temperature. Cells were washed twice with distilled water and observed using a Invitrogen EVOS M7000 Cell Imaging System (ThermoFisher Scientific) with a 4x objective.

### Histological Analysis

Interscapular brown adipose and inguinal white adipose tissues were harvested and fixed in 10% formalin for 24 hours. Fixed tissues were dehydrated in increasing concentrations of ethanol, cleared with xylene, and embedded in paraffin. Paraffin blocks were cross-sectioned (5 μm), and sections were then deparaffinized and stained with hematoxylin-eosin. Slides were observed using an Invitrogen EVOS XL Core Imaging System (ThermoFisher Scientific). Images presented were taken with a 40x objective.

### Western Blotting

For tissue samples, 50-100 mg of tissue were mechanically lysed in extraction buffer (50 mM Tris HCl, 5 mM EDTA, 200 mM NaCl, pH = 7.4) supplemented with protease and phosphatase inhibitors (Sigma, USA). Samples were centrifuged to remove excess fat and debris and incubated with chemical detergents as previously described (45). Cell samples were lysed in RIPA buffer (Boston Bioproducts, USA) supplemented with protease inhibitors (Sigma, USA) and centrifuged at 12.000 rpm at 4°C for 5 min. Protein concentration was determined through the BCA method and samples were diluted in Laemli buffer and separated through precast 8-12% polyacrylamide denaturing gels (BioRad, USA), in Tris-glycine buffer. Samples were transferred to nitrocellulose and PVDF membranes and stained with Ponceau. Membranes were blocked in 5% milk in Tris-buffered saline with 0.1% Tween 20 detergent (TBST) and incubated with primary antibodies diluted in 5% BSA solution overnight at 4°C. After incubation, membranes were washed in TBST, and horseradish peroxidase (HRP)-conjugated secondary antibodies anti-Rabbit (7074S) were added. SuperSignal West Femto (ThermoFischer Scientific) was used as an HRP substrate. Band detection was performed using ChemiDoc equipment (BioRad Laboratories, USA). Alternatively, fluorescent secondary antibodies (IRDye® anti-mouse and anti-Rabbit) were added, and band detection was also performed using ChemiDoc.

### qPCR

Cells were lysed using TRIzol® reactant followed by RNA phase extraction and precipitation (46). 0.6 to 1 μg of RNA were employed for cDNA synthesis (High-Capacity System, Life Technologies, USA). Real-time RT-PCR analysis of gene expression was performed using a QuantStudio 6 detection system (Life Technologies, USA). Each PCR reaction contained 10 ng of cDNA, 200 nM of each specific primer, PowerUp SYBR Green PCR Mastermix (Life Technologies, USA), and nuclease-free water to a final volume of 10-20 μL. Cyclophilin gene expression was used as endogenous control.

### Seahorse Extracellular Flux Analysis

Seahorse XFe24 and XFe96 were employed to assess cellular oxygen consumption and extracellular acidification rates. For preadipocyte assessments, 1000 cells/mm^2^ were seeded on a Seahorse cell culture plate one day before the experiment. For differentiated adipocytes, 1500 preadipocyte cells/mm^2^ were seeded, and after 2 days, adipocyte differentiation was induced and carried out as described. Before experiments, cells were washed and incubated in serum-free experimental culture media (high glucose DMEM, containing 1 mM pyruvic acid and 2 mM glutamine) for 1 h at 37°C. Oxygen consumption measurements were performed at baseline conditions, in the presence of oligomycin (3 μM), followed by FCCP stimuli (20 μM) and inhibition by antimycin A (5 μM) plus rotenone (5 μM). Ideal quantities of inhibitors were titrated in preliminary experiments (not shown). Alternatively, cells were treated with equal volumes of dilutant DMSO, diazoxide (60 μM), or diazoxide plus glibenclamide (5 μM). Adipocytes were then stimulated with norepinephrine (50 μM), followed by oligomycin (3 μM), and an antimycin A (5 μM) plus rotenone (5 μM) combination.

### Resipher Detection of Cultured Cell Oxygen Consumption

The Resipher system (Lucid Scientific, USA) continuously and non-invasively monitors oxygen consumption in 96-well plates by measuring oxygen gradients within wells. WT-1 preadipocytes were seeded on commercial 96-well plates at 1500 cells/mm^2^. After one day of acclimatation, preadipocyte oxygen consumption was detected by placing the Resipher sensors over the cell culture plate, following manufacturer instructions. Adipocyte differentiation was then carried out as described above.

### Mitochondrial Isolation

Mitochondria from BAT were isolated as described previously, with slight modifications (4). Interscapular (iBAT) collected from two to three animals was washed in cold PBS and suspended in 5% (w/v) isolation buffer (240 mM sucrose, 5 mM Hepes, 1% fatty acid free BSA, pH = 7.2). The suspension was homogenized using an Elvehjem manual potter homogenizer. The homogenate was filtered and centrifuged at 8500 g for 10 min, at 4°C. The fat layer and supernatant were discarded, and the pellet was suspended in the same volume of isolation buffer, transferred to a new tube, and centrifuged at 700 g for 5 min at 4°C, twice. After each centrifugation, the supernatant was transferred to a new tube and the pellet was discarded. Following this, the supernatant was submitted to another centrifugation at 8500 g. The pellet was washed in suspension buffer (240 mM sucrose, 5mM Hepes, pH = 7.2). After a final centrifugation at 8500 g for 10 min, mitochondria were suspended in a minimal volume of suspension buffer, and protein concentration was determined using the Bradford method (47).

### Mitochondrial Swelling

Mitochondrial swelling was assessed through the light scattering technique, as described previously (15, 48), and based on the lower light scattering associated with more swollen matrixes. Briefly, 35 μg of mitochondrial protein were suspended in a hypotonic experimental buffer (100 mM KCl, 10 mM Hepes, 2 mM KH_2_PO_3_, 2 mM MgCl_2_, 2 mM succinate, with or without 2 mM pyruvate and malate, pH = 7.4), while light scattering was measured at 520 nm and 90° angle using a 4500 Hitachi fluorimeter, at 37°C under constant stirring. Succinate (2 mM) and GDP (2 mM) were added to the experimental buffer prior to mitochondrial addition.

### Oxygen Consumption by Isolated Mitochondria

In a high-resolution O2k respirometer (Oroboros, Australia), 50 μg of mitochondrial protein were suspended in respiration buffer (125 mM sucrose, with 65 mM KCl, 10 mM Hepes, 2 mM phosphate, 2 mM MgCl_2_ and 0.2% bovine serum albumin, adjusted to pH 7.2 with KOH) under constant stirring. Antimycin A (2 μM) and TMPD (2 mM) plus ascorbate (2 mM) were added. GDP (2 mM) was used to inhibit UCP1, while oligomycin (2 μM) was used to inhibit ATP synthase. Diazoxide (60 μM) or a combination of diazoxide plus glibenclamide (5 μM) were added from beginning to activate or inhibit MitoK_ATP_, respectively.

### Statistical Analysis

All data were analyzed using GraphPad Prism 7.0 (GraphPad Software, USA) and represented as averages ± SEM. Statistical analyses were conducted through t-tests, one-way or two-way ANOVA followed by Holm-Sidak’s post-hoc test, when appropriate.

## Results

### MitoKATP protein levels change in response to BAT activation

Based on our hypothesis that MitoKATP could be of particular importance in BAT mitochondria due to their low inner membrane potentials upon activation, we checked the GeneAtlas MOE430 gcrma dataset (49) using BioGPS (50, 51), which profiles gene expression from a diverse array of normal mouse tissues, organs, and cell lines. We found that both genes annotated to form the MitoKATP holo-protein, Ccdc51 (MitoK), coding for the pore-forming subunit, and Abcb8 (MitoSur), coding for the regulatory portion (19), are expressed in mouse BAT. Interestingly, MitoSur is expressed at particularly high levels in BAT when compared to all other mouse tissues: approximately 4 times higher than in white adipose tissue and skeletal muscle and double the expression levels of liver and kidney.

Next, we aimed to detect the peptide products of both genes in mouse BAT after 14 days exposure to thermoneutrality or cold (28). As expected, cold exposure promoted significant tissue remodeling relative to thermoneutrality conditions, including lower triglyceride content (Fig. 1a) and higher UCP1 levels (Fig. 1b, d), while mitochondrial components Timm23 (Fig. 1e) and oxidative phosphorylation components (Fig. 1f) remained unchanged. Interestingly, MitoK levels in cold-exposed BAT presented high heterogeneity relative to thermoneutral conditions, with a trend towards a slight upregulation (Fig. 1c). We were unable to detect MitoSUR levels reliably in these mouse samples.

**Figure 1.**
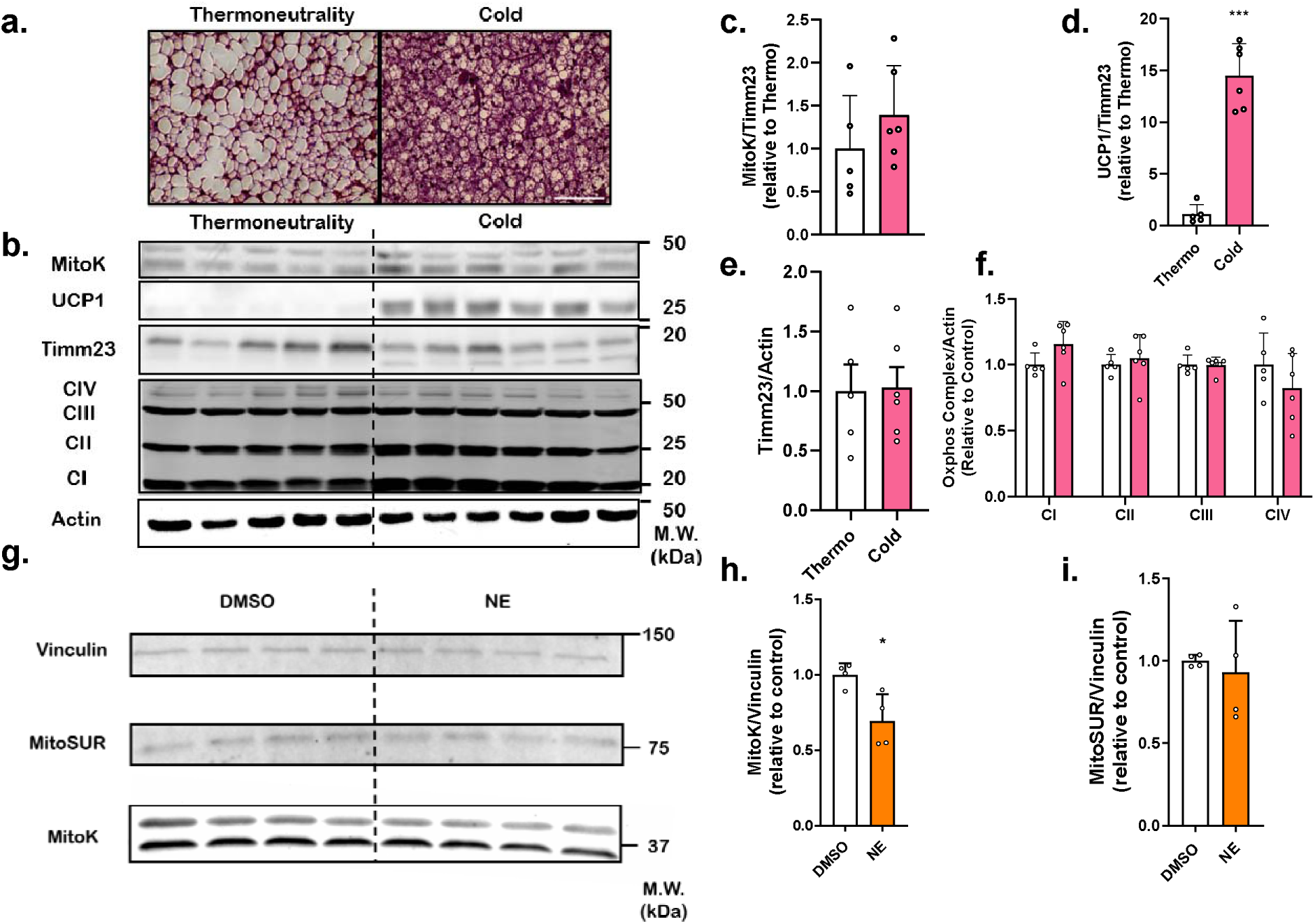
MitoKATP components are expressed in BAT and respond to acute stimulation. (a) HE-stained BAT sections from mice maintained in thermoneutrality or cold for 14 days, revealing clear tissue remodeling at different temperatures. Scale bar, 100 μm. (b) MitoK, UCP1, Timm 23, and oxidative phosphorylation component (total OxPhos) protein levels estimated by western blotting, relative to the actin loading control. (c-f) Quantifications of normalized band densitometries from panel b. (g) Differentiated 9B cells were treated with norepinephrine (NE) or an equivalent amount of dilutant DMSO for 24 h. Cells were then blotted for MitoK, MitoSUR, and Vinculin as a loading control. (h, i) Quantifications of normalized band densitometries from panel e. * p < 0.05, *** p < 0.001, Student’s t-test.

The effects of BAT activation on the expression of MitoKATP components were also studied in a more acute and *in vitro* model, which presents higher homogeneity. The 9B cell line of immortalized mouse preadipocytes was differentiated towards mature adipocytes. Cells were then stimulated with norepinephrine, and protein levels were assessed 24 hours later (Fig. 1g). We found that stimulated cells presented a decrease in MitoK protein levels (Fig. 1h), but unchanged quantities of MitoSUR (Fig. 1i). Overall, these data indicate that changes in BAT activity seem to promote changes in the amounts of protein components of the MitoKATP.

### Genetic ablation of MitoKATP hampers proliferation and metabolic fluxes in human pre-adipocytes

To verify the role of MitoKATP in BAT, we attempted to produce knockout cells, initially employing the A41hBAT cell line, an immortalized preadipocyte cell line originated from human cervical BAT, which has the advantage that it can fully differentiate into functional, UCP1 expressing, brown adipocytes (37). For MitoK downregulation, we generated stable cell lines expressing Cas9, and a single gRNA targeting either MitoK or no target against the human and mouse genomes, as our control.

Our strategy was efficient in downregulating the MitoK gene through two different gRNA sequences, compared to our non-targeted gRNA (gNT) (Fig. 2a). Notably, MitoK-deficient cells displayed decreased proliferation (Fig. 2b), as evidenced by crystal violet staining. Control A41hBAT preadipocytes reached high confluency after being seeded at low density, whereas mutant adipocytes proliferated notably less within six days.

**Figure 2.**
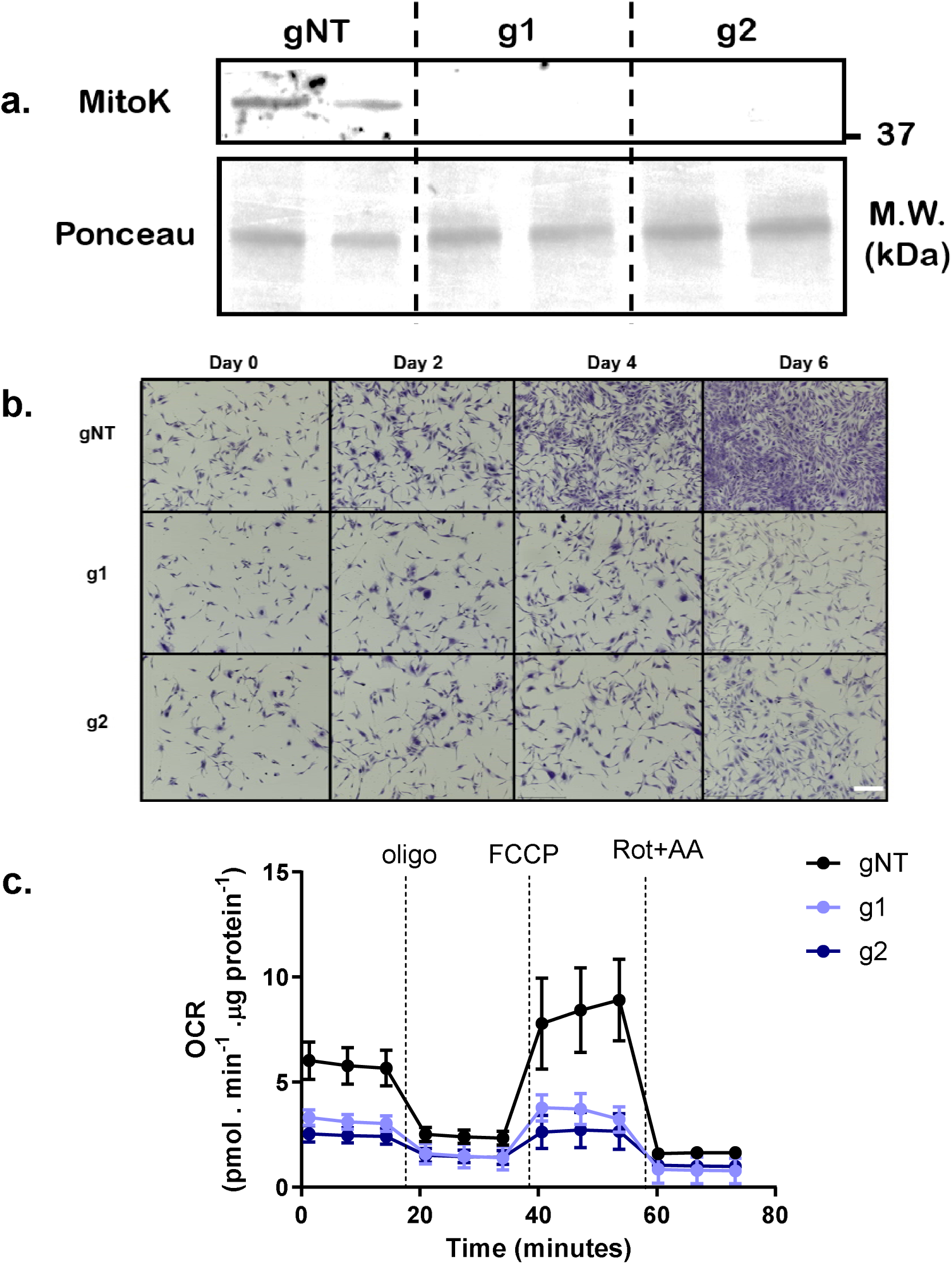
CRISPR-induced MitoK deletion in preadipocytes hampers proliferation and differentiation. (a) A41hBAT preadipocytes presented effective deletion of MitoK relative to non-targeted gRNA (gNT). (b) A41hBAT cells were seeded at the same density at day 0 and fixed on days 0, 2, 4, or 6. Scale bar, 400 μm. (c) Mitochondrial oxygen consumption was assessed in Seahorse XFe experiments, under basal conditions and after the addition of ATP-synthase inhibitor oligomycin (oligo), uncoupler FCCP and electron transport inhibitors rotenone plus antimycin (Rot+AA), as described in the methods section.

The proliferative deficiency was associated with a marked decrease in mitochondrial oxygen consumption in MitoK deficient preadipocytes (Fig. 2c), including a strong decrease in oligomycin-sensitive respiration, or ATP-linked respiration, as well as decreased FCCP-stimulated oxygen consumption rates, or maximal electron transport capacity. No changes in non-mitochondrial respiration (in the presence of rotenone plus antimycin) were registered. These results indicate that MitoK is important to maintain mitochondrial electron transport, oxidative phosphorylation, and proliferation in human brown adipocytes. Unfortunately, these results, while interesting, also indicate that this cell line cannot be used to evaluate the effects of MitoKATP in mature adipocytes, given that proliferation of preadipocytes is essential for adipocyte differentiation (38).

### Genetic ablation of MitoKATP affects mouse preadipocyte metabolism

A second strategy involved the use of the mouse preadipocyte cell line WT-1 (35). Again, we achieved effective downregulation of MitoK protein levels (Fig. 3a), now without associated proliferative deficiency (not shown). Similarly to the human cell line, preadipocytes deleted for MitoK exhibited lower ATP-linked (oligomycin-sensitive) and maximal (FCCP-stimulated) oxygen consumption rates (Fig. 3b). Additionally, as an alternative strategy to downregulate the MitoK gene, shRNA expression was used, and consistently led to decreased oxygen consumption levels (Sup. Fig. 1). This demonstrates that MitoK is necessary for the maintenance of electron transport capacity in preadipocytes.

**Figure 3.**
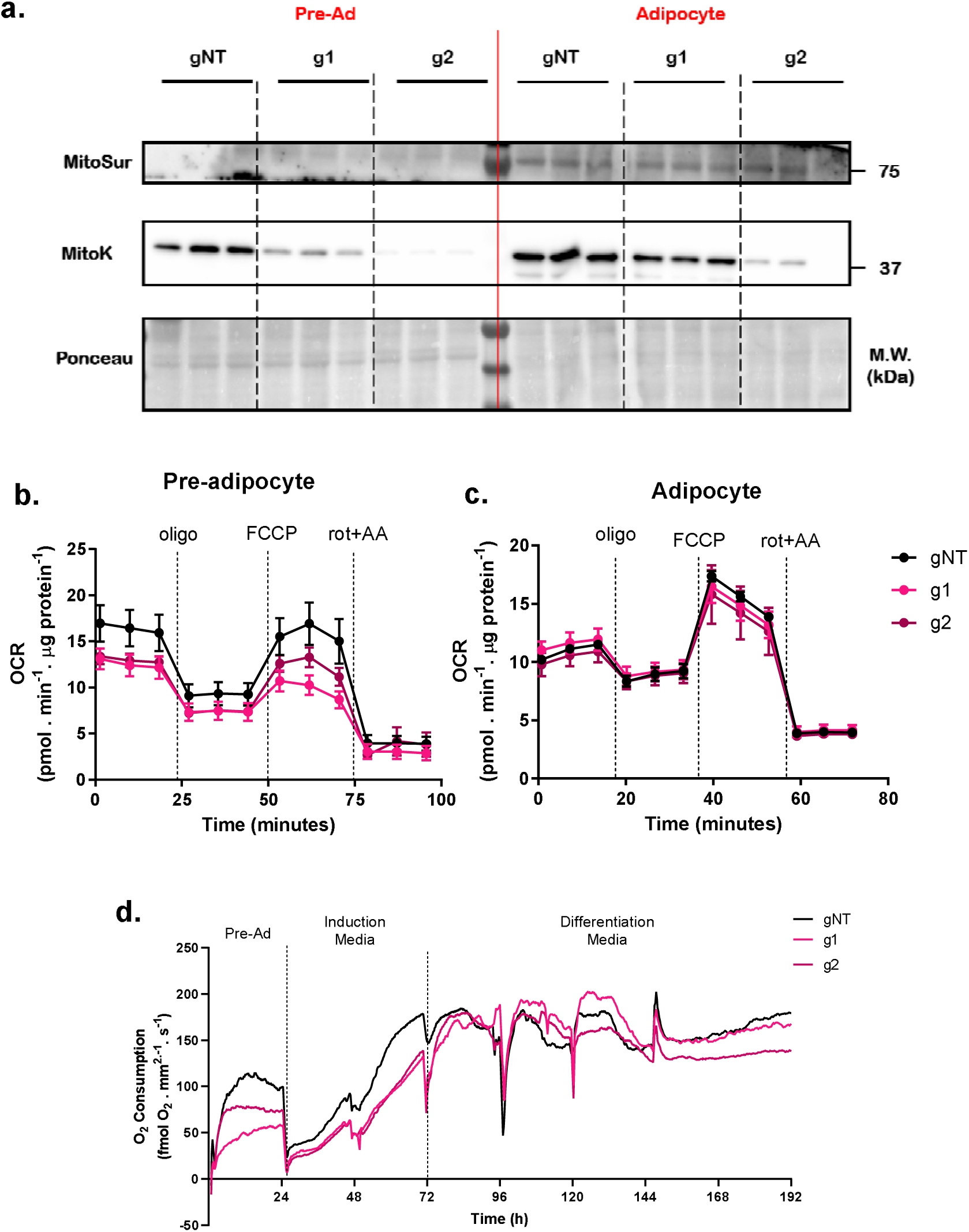
MitoK-deficient preadipocytes, but not mature adipocytes, present decreased oxygen consumption rates. (a) WT-1 preadipocytes presented effective decrease of MitoK relative to non-targeted gRNA (gNT). This pattern was maintained during the mature adipocyte stage. (b) MitoK-deficient cell lines displayed decreased oxygen consumption in the preadipocyte stage, but not in the mature adipocyte stage (c). (d) Oxygen consumption rates are lower in MitoK-deficient cells during the preadipocyte and early induction stages, normalizing relative to gNT after 72 h.

Since the MitoK-deficient WT-1 cells reached confluency similarly to their control counterparts, we submitted preadipocytes to differentiation protocols and confirmed that cells could upregulate a series of gene markers of adipogenesis, as well as brown fat markers (Sup. Fig. 2). Importantly, although adipocyte differentiation promotes an upregulation in the MitoK gene, the pattern of gene downregulation observed in the preadipocyte stage was maintained for differentiated cells (Fig. 3a). Although some gene downregulation was maintained, the previously observed decrease in oxygen consumption was no longer present in fully differentiated adipocytes (Fig. 3c).

In order to understand the dynamic change in metabolic fluxes during differentiation, we employed the Resipher system, capable of measuring oxygen consumption of cultured cells throughout several days under normal cell culture conditions. We saw that mutant cells displayed decreased cellular oxygen consumption rates during the preadipocyte stage (Fig. 3d). However, this difference occurred during differentiation up to 72 h. After this time point, immature mutant brown adipocytes already exhibited oxygen consumption levels comparable to those of control gNT cells (Fig. 3d). Thus, our results indicate that MitoKATP plays different roles in preadipocyte and mature brown adipocyte metabolism, and that adipocyte differentiation may trigger compensatory mechanisms for the lack of MitoKATP activity, recovering oxidative phosphorylation capabilities in mature brown adipocytes. Alternatively, our data also suggest that MitoKATP, while clearly modulating oxygen consumption in preadipocytes, may not play a role in oxidative phosphorylation in mature brown adipocytes.

### Acute inhibition of MitoKATP leads to increased adrenergic-induced oxygen consumption in mature adipocytes

Since genetic ablation of MitoK changed metabolic parameters in preadipocytes, which do not respond to adrenergic stimuli, and promoted metabolic adaptations during differentiation, we sought to investigate the effects of MitoKATP on mature brown adipocytes through acute activation and inhibition using pharmacological modulators diazoxide (a MitoKATP activator, MitoKa) and glibenclamide (a MitoKATP inhibitor, MitoKi). Cells were grown under equal conditions, and therefore did not present significant changes in basal respiration (results not shown), so data were normalized to basal levels. Addition of MitoKa or MitoKi did not change basal respiration significantly. Using primary brown adipocytes (Fig. 4a), a large increase in oxygen consumption rates can be induced by the addition of norepinephrine (NE). As expected for BAT cells, this oxygen consumption is uncoupled to ATP production as seen by the lack of effect of added ATP synthase inhibitor oligomycin (oligo), but is due to mitochondrial electron transport, as seen by inhibition induced by antimycin A plus rotenone (AA + rot). Interestingly, the inhibition of MitoKATP (MitoKi) significantly increased NE-induced brown adipocyte activation in these cells, while the addition of an activator did not significantly change NE-induced oxygen consumption (quantified in Fig. 4c), indicating that maximal adrenergic responses requires MitoKATP inhibition. While these results are surprising, they were also observed using cell-line derived differentiated brown adipocytes (Fig. 4b, d).

**Figure 4.**
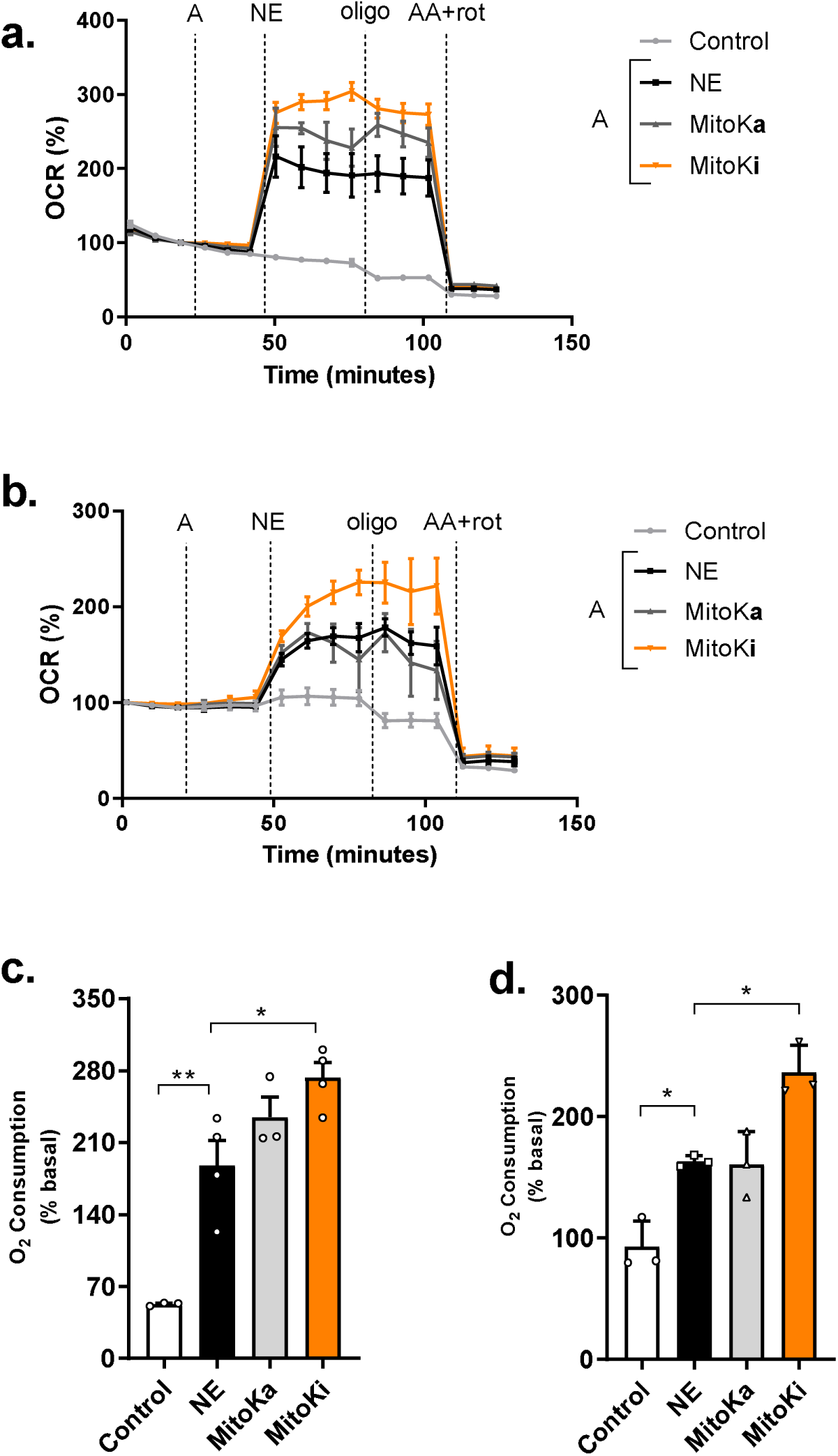
Full activation of brown adipocytes requires MitoKATP inhibition. Oxygen consumption rates were assessed in primary brown adipocytes (a, c) or differentiated 9B cells (b, d) treated with pharmacological MitoKATP activator (MitoKa) or inhibitors (MitoKi). NE, oligomycin and AA + rot were added as indicated. * p≤ 0.05, ** p≤ 0.01 one-way ANOVA, with Holm-Šidak’s post-hoc test.

Overall, these results indicate that MitoKATP is largely open in untreated brown adipocytes (leading to no response to pharmacological activators), but these cells are more strongly activated, promoting more pronounced uncoupled respiration, when MitoKATP is pharmacologically inhibited.

### Decreased MitoKATP activity enhances UCP1-driven oxygen consumption

Given that NE-stimulated brown adipocytes presented enhanced uncoupling activity in vitro when MitoKATP was inhibited, we sought to see if the same was true for primary brown adipose tissue *in vivo*. To do so, we employed CL316,243, a selective adrenergic agonist capable of stimulating thermogenic fat depots (30, 31) without systemic adrenergic effects. Animals were injected daily for two weeks, and compared to saline-treated counterparts. As expected, CL316,243-treated animals had lower weight gain, accompanied by a clear remodeling of interscapular brown and inguinal fat depots, indicating effective adrenergic agonism in different fat pads (Sup. Fig. 3).

After the two week treatment period, we analyzed the brown fat depots and confirmed that CL316,243 increased UCP1 content in isolated mitochondria from BAT (Fig. 5a, b). Interestingly, and similarly to what we had previously seen with cold exposure, we didn’t observe overt changes in the protein content of MitoK in these mitochondria (Fig. 5c-e). However, the quantity of the protein does not indicate activity. Indeed, MitoKATP has been extensively described to be modulated by endogenous signals and cellular states (16, 26, 52, 53). To assess MitoKATP activity, we employed swelling assays (Fig. 5f), which are based on the premise that isolated mitochondria are K^+^-depleted and present matrix shrinkage. As a result, suspension in K^+^-rich experimental media after isolation leads to changes in mitochondrial volume, associated with K^+^ uptake and osmotically obligated water entry, resulting in decreased light scattering over time. Due to specific characteristics of BAT, these experiments had to be designed to maintain MitoKATP open while minimizing the effects of UCP1-mediated uncoupling, in order to achieve measurable K^+^ uptake and swelling. This was achieved by incubating mitochondria with GDP, which opens MitoKATP and inhibits UCP1 (although full UCP1 inhibition also requires albumin, which cannot be used in swelling experiments due to its oncotic effects). Under these conditions, we observed that mitochondria from CL316,243-treated animals swell at slower rates than mitochondria from saline-treated animals, indicating decreased K^+^ import rates (Fig. 5f) and inhibited MitoKATP activity.

**Figure 5.**
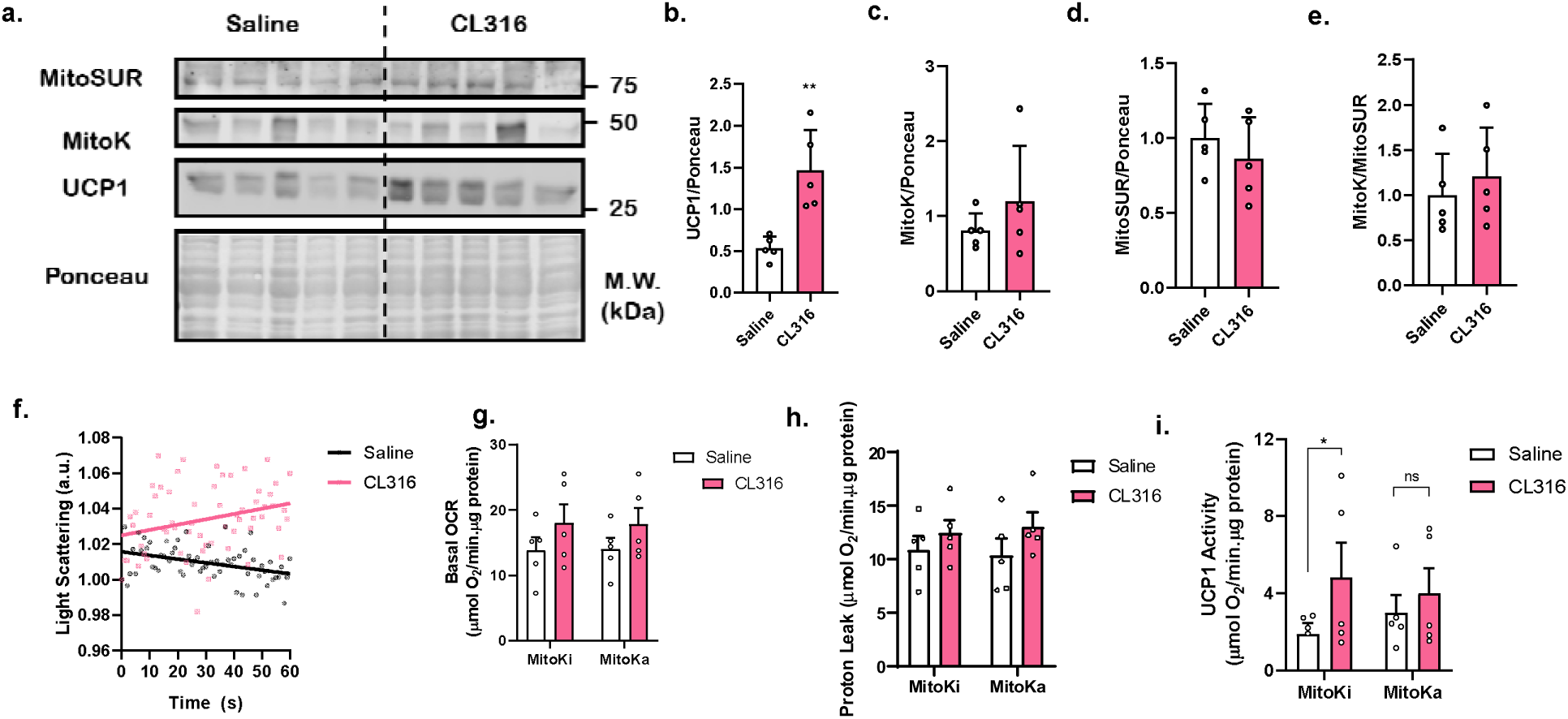
MitoKATP inhibition is required for full uncoupling activity in stimulated BAT mitochondria. (a-e) Mitochondria were isolated from BAT obtained from CL316-treated animals, and blotted for UCP1, MitoK, and MitoSUR. (f) BAT mitochondrial swelling associated with K^+^ uptake was assessed in mitochondria isolated from control or CL316-treated animals, in the presence of GDP to activate MitoKATP, while maintaining UCP1 inhibited. (g) basal OCR were assessed in isolated mitochondria from control or CL316-treated animals, in the presence of MitoKATP inhibitors (MitoKI) or activators (MitoKA), as described in the Methods section. (h) proton leak-associated OCRs were measured after additions of 2 mM GDP and 2 μM oligomycin. (i) UCP1 activity was estimated in isolated mitochondria by inhibiting the protein with 2 mM GDP. * p < 0.05, T-test panels b-g, Two-Way ANOVA panels h and i.

Given our results in BAT cells, in which full activation of UCP1-mediated uncoupling was only achieved when MitoKATP was inhibited, we questioned if the change in activity of MitoKATP observed in CL316,243-treated animals participated in the activation of uncoupling in these mitochondria. We measured basal (Fig. 5g), oligomycin- and GDP-insensitive proton-leak (Fig. 5h), and UCP1-associated (Fig. 5i) oxygen consumption in these mitochondria, as promoted by GDP-mediated UCP1 inhibition, in the presence of MitoKa or MitoKi. All mitochondrial preparations had similar basal and GDP-insensitive proton leaks (Fig. 5g,h). This demonstrates that CL316,243 treatment does not significantly change lipid composition and proton permeability, and that the activity of MitoKATP in BAT does not promote measurable uncoupling, as seen in other tissues (15). UCP1-associated oxygen consumption was, as expected, enhanced in mitochondria from CL316,243-treated animals (Fig. 5i). However, these differences are lost when MitoKATP is activated (MitoKa), confirming our prior results indicating that MitoKATP inhibition is important to promote full BAT UCP1-mediated uncoupling.

## Discussion

Here, we studied the role of MitoKATP channels in brown adipose tissue differentiation and function. Prior to this work, there were no studies regarding MitoKATP channels in this tissue, as assessed by probing publication databases for papers relating to BAT, MitoKATP, MitoK, or MitoSUR. This is a significant gap, given that mitochondria in BAT exhibit many different characteristics to other tissues which would be expected to alter and modulate K^+^ transport in these organelles.

K^+^ entry into mitochondria is expected to occur through K^+^ leakage into the matrix, attracted by the negative matrix charge, and facilitated by the high concentrations of the ion in the cytosol (14). Since BAT mitochondria have lower inner membrane potentials, this could result in lower K^+^ transport into the matrix, an effect that we hypothesized could be compensated by higher MitoKATP activation in this tissue.

More evidence that K^+^ homeostasis is particularly important in BAT mitochondria comes from early studies establishing conditions to study isolated organelles from this tissue. While most isolated sucrose buffer-stored mitochondria lose matrix water and re-equilibrate their matrix volume within the first few seconds in experimental media (14–16), mitochondria isolated from BAT present an unusually highly condensed matrix and must be in low osmolarity media to expand the matrix to levels similar to those seen *in vivo*. This incubation condition is essential to obtain full respiratory capacity *in vitro* (4). Alternatively also, BAT mitochondria could recover respiration in isosmotic intracellular media over a period of several minutes, as their matrixes expanded (54).

Together, these results suggested that BAT mitochondria had particular K^+^ transport properties dissimilar to other tissues. Indeed, we found that GeneAtlas data indicate that MitoKATP components are highly expressed in BAT, suggesting this channel is specifically important for this tissue. This is of interest since MitoKATP components are calculated to be of low abundance in general, in the region of 1 fmol or less of channel per mg of mitochondrial protein (55). Based on this, we hypothesized that MitoKATP could play an important role in BAT function. Indeed, supporting this idea, we found changes in the levels of MitoKATP components in animals and cells treated under conditions that change BAT activity (cold exposure and NE-stimulation; Fig. 1).

While attempting to produce MitoKATP-ablated BAT cells, we found that loss of MitoKATP profoundly decreased mitochondrial electron transport capacity in preadipocytes (Fig. 2 and 3), to the point in which human cell lines were functionally hampered and unable to differentiate without expression of the channel (Fig. 2). In cells in which the absence of MitoKATP did not impede differentiation, the respiratory inhibition promoted by MitoKATP deficiency persisted during the preadipocyte stage, but was not seen in mature adipocytes (Fig. 3). Together, these results demonstrate that MitoKATP is important to maintain electron transport capacity during BAT differentiation.

In mature adipocytes, we expected that MitoKATP could be important to maintain matrix volumes under conditions in which UCP1 was activated, leading to uncoupling. To our surprise, instead, we found quite the opposite: MitoKATP pharmacological activation did not significantly change UCP1-mediated NE-stimulated uncoupled oxygen consumption in two different BAT cell lines. Conversely, MitoKATP inhibition strongly enhanced mitochondrial uncoupling in both cell types (Fig. 4), indicating that the channel must be fully inactive to promote full uncoupling.

Further supporting this notion, mitochondria isolated from mice treated with a selective BAT adrenergic agonist exhibited lower matrix swelling rates compatible with lower MitoKATP activity (Fig. 5). Maximal mitochondrial uncoupling was seen in these mitochondria only when MitoKATP was fully inhibited, while the activity of the channel did not lead to changes in UCP1-independent proton leak. These data suggest that pharmacological UCP1 activation *in vivo* is accompanied by MitoKATP inhibition, which is essential to support ideal UCP1-mediated uncoupling activity.

Overall, our findings are the first to indicate that MitoKATP is an important regulator of both BAT differentiation and thermogenic function, suggesting that K^+^ homeostasis should be further studied in this tissue, as a complementary and functionally interconnected activity to uncoupling in brown adipocytes.

**Supplementary figures and legends:** doi.org/10.5281/zenodo.15577735

## Supporting information

supl mat

## Acknowledgements

The authors thank Silvânia M. P. Neves and her animal facility crew for exceptional expert animal care. This work was supported mainly by the Fundação de Amparo à Pesquisa do Estado de São Paulo (FAPESP) under grant numbers 13/07937-8, 20/06970-5 and 20/14159-5 as well as the Conselho Nacional de Desenvolvimento Científico e Tecnológico (CNPq) and Coordenação de Aperfeiçoamento de Pessoal de Nível Superior (CAPES) line 001.

## Notes

### Competing Interest Statement

The authors have declared no competing interest.

### Summary of Updates

New experiments, new figures, new discussion.

