## Supplementary material for "Mitochondrial ATP-Sensitive K^+^ Channels (MitoK_ATP_) Regulate Brown Adipocyte Differentiation and Metabolism": supl mat

### Supplementary Figures

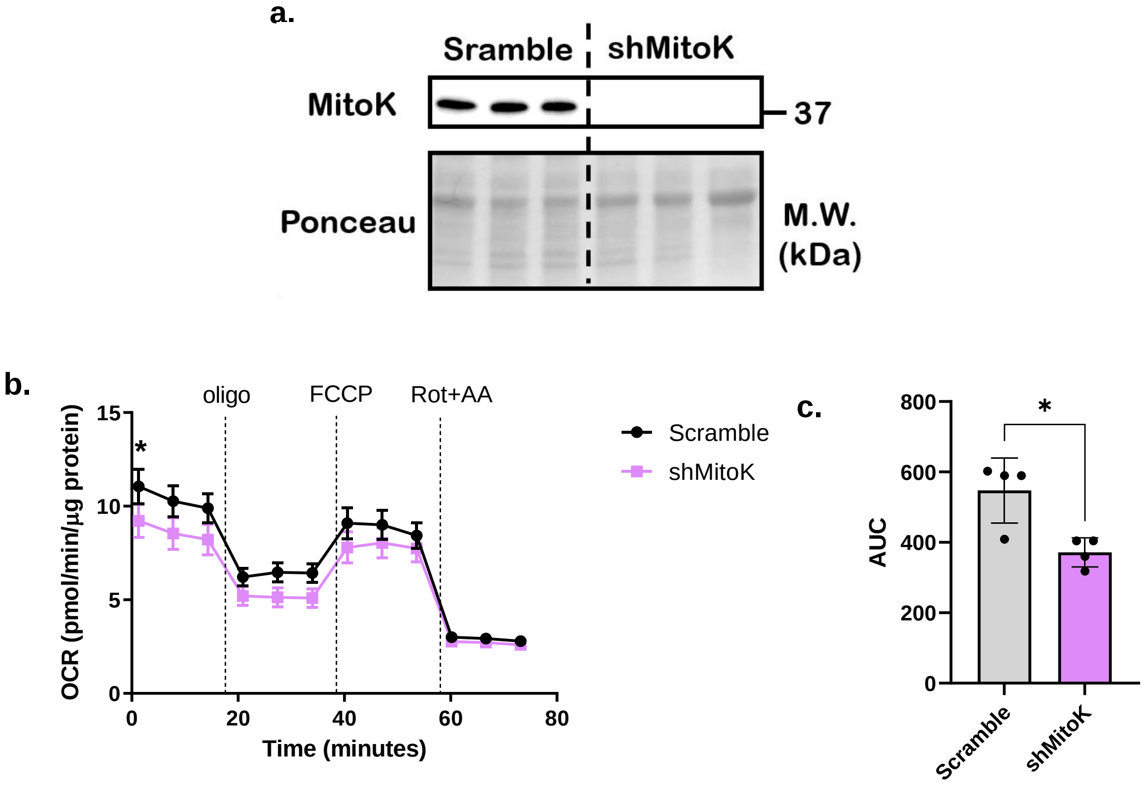


**Sup. Fig. 1. MitoK KD preadipocytes have deficient oxygen consumption.** shRNA delivery was validated by decreased MitoK protein levels (a). Decreased levels of MitoK correlated with impaired oxygen consumption (b). The total area under the curve was calculated from the respirometry experiments (c). * $p < 0.05$, paired t-test.

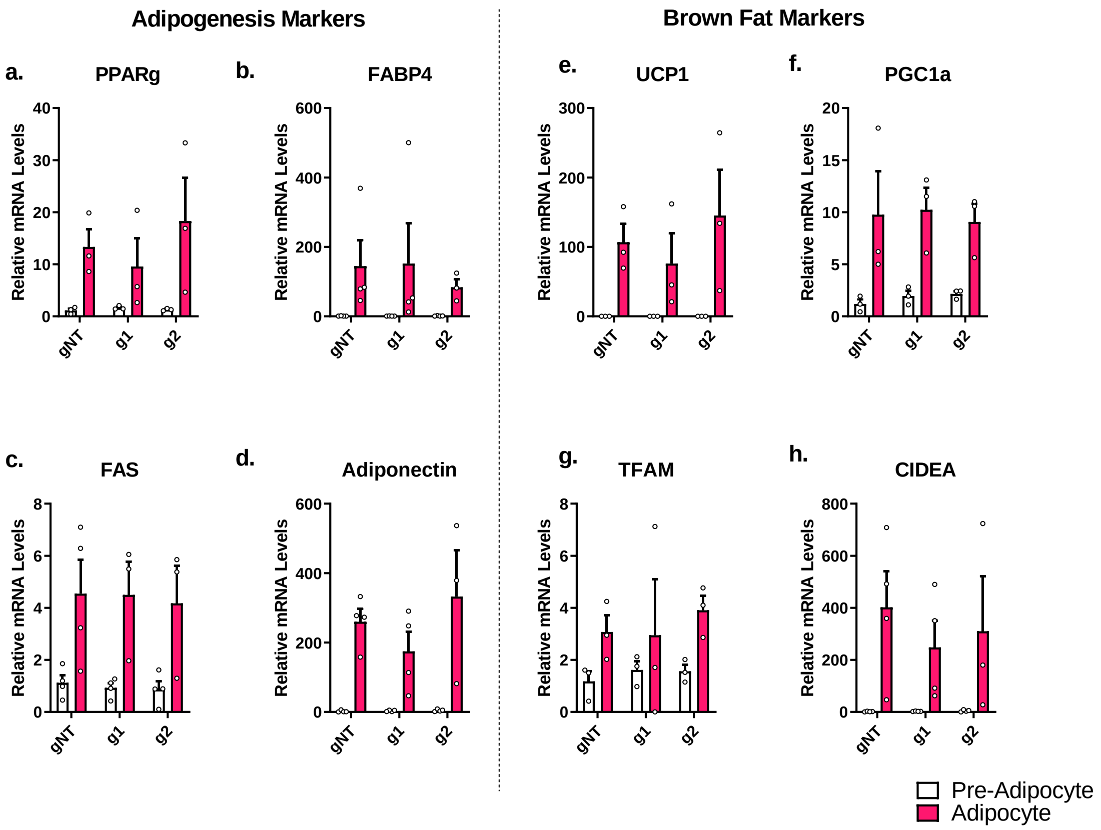


**Sup. Fig. 2. WT-1 MitoK mutant preadipocytes differentiate into mature adipocytes similarly to control preadipocytes.** WT-1 cell lines were submitted to a differentiation protocol. By day 8, cells were collected for relative mRNA quantification. Pparg (a), FABP4 (b), FAS (c) Adiponectin (d), UCP1 (e), PGC1a (f), TFAM (g) and CIDEA (h) were all upregulated by adipocyte differentiation. No significant changes were observed when comparing mutant groups with gNT cells.

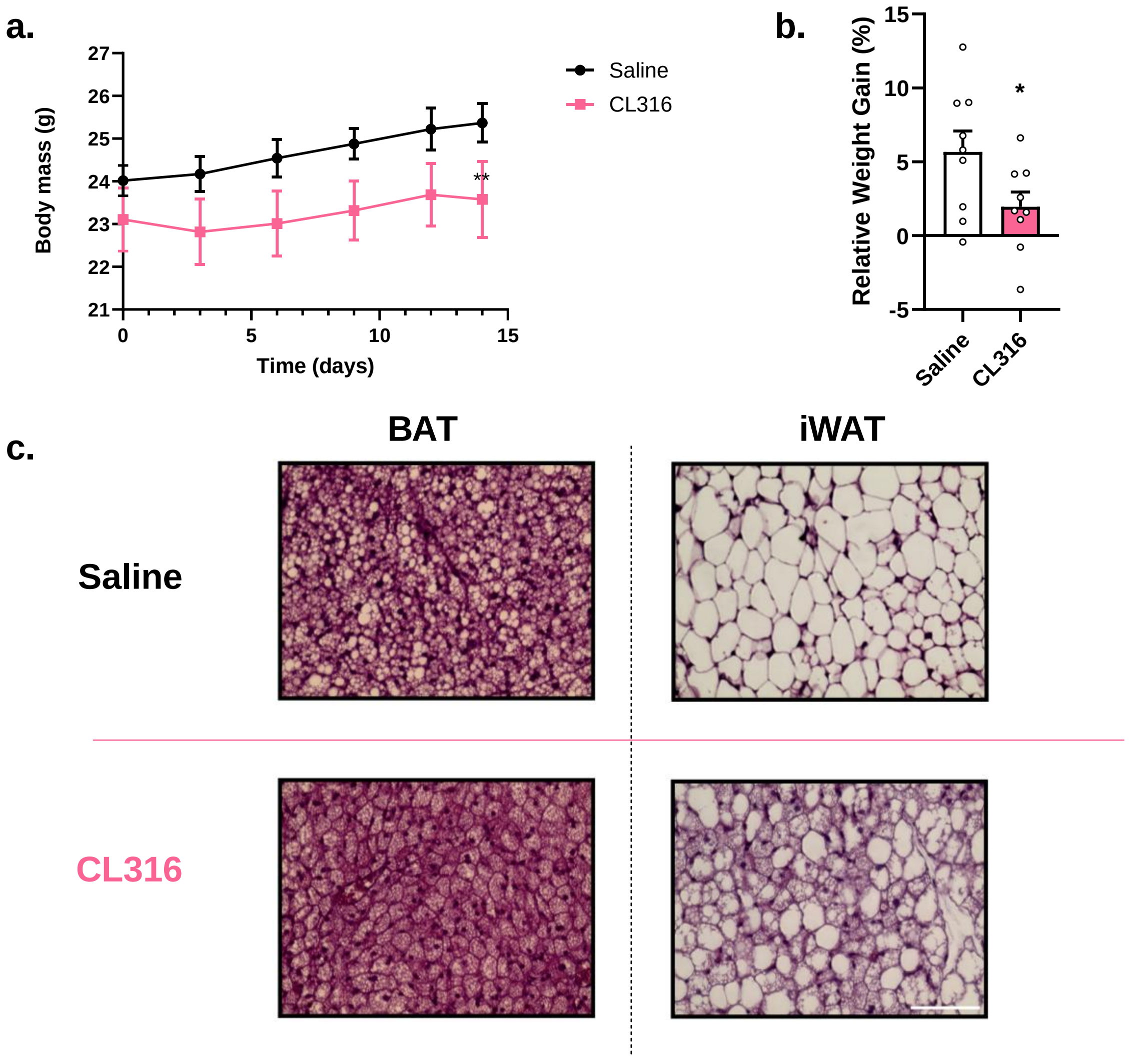


**Sup. Fig. 3. CL316,243 treatment affects body weight and BAT.** Animal body weight was assessed every other day throughout CL-316 treatment (a). CL316-treated animals have decreased weight gain, when compared to saline-injected mice (b). HE staining of the BAT and inguinal WAT shows decreased lipid droplet area and emergence of multilocular adipocytes in the white fat pad. Scale bar, 100 μm (c).
